# The gut microbiota-derived metabolite queuosine regulates neuronal development and network function through tRNA modification

**DOI:** 10.64898/2026.07.02.736043

**Authors:** Ning Yang, Yu Sun, Lisa Mallis, Marin Boutonnet, Julia Bär, Ann E. Ehrenhofer-Murray, Marina Mikhaylova

## Abstract

Queuosine (Q) modification is a hypermodified nucleoside derived from guanine on tRNAs that enhances the decoding of codons and equilibrates translational speed. Q is biosynthesized in bacteria, and eukaryotes salvage Q and the nucleobase queuine from the diet and the gut microbiome. In animals, Q deficiency causes impaired proteostasis, mitochondrial dysfunction, and neurological phenotypes, possibly due to the longevity and high metabolic demand of neurons. Yet, how Q affects isolated neurons has not been explored yet. Here, primary rat cortical neurons were cultured in Q-free synthetic medium to directly modulate Q modification levels independently of genetic perturbation, enabling assessment of its effects on neuronal development, survival, morphology, synaptic organization, and activity. Importantly, we found that the presence of Q modification facilitated neuronal arborization, decreased inhibitory synaptic density, and increased the frequency of spontaneous calcium transients, showing that tRNA Q modification enhances neuronal structural maturation and synaptic activity. Thus, the fine-tuning of neuronal translation programs by Q-tRNAs is required for proper network development and may influence neuronal resilience and synaptic function.

## Introduction

Queuosine (Q) modification is found at the wobble position 34 of tRNAs with a GUN anticodon, which decode asparagine, aspartic acid, histidine, and tyrosine codons (Ehrenhofer-Murray, 2025) (Figure 1A). Q is a 7-deazaguanosine derivative with a cyclopentene diol moiety that preserves the canonical G-like Watson-Crick pairing with C but improves non-canonical wobble base pairing with U (Huber *et al*, 2023). Although Q modification is conserved in eukaryotes and bacteria, the complete biosynthetic pathway for Q from GTP is only present in bacteria (de Crecy-Lagard *et al*, 2024). Eukaryotes cannot synthesize Q and therefore rely on dietary sources and the microbiota to obtain the Q nucleoside and its nucleobase queuine (q). Both Q and q are transported into the cell by the transmembrane transporter SLC35F2/ Qtp1 (Burtnyak *et al*, 2025), and Q is hydrolyzed to release the q base (Hung *et al*, 2023; Patel *et al*, 2022). Queuine is the substrate for the tRNA guanine transglycosylase enzyme (TGT) that replaces G34 by q in the respective tRNAs (Chen *et al*, 2010). Eukaryotic TGT is a heterodimeric enzyme, consisting of the subunits QTRT1 and QTRT2 that are both homologs of bacterial TGT. In eukaryotes, tRNA^Asp^ and tRNA^Tyr^ can be further glycosylated with mannose and galactose, respectively, to yield manQ and galQ (Zhao *et al*, 2023).

**Figure 1.**
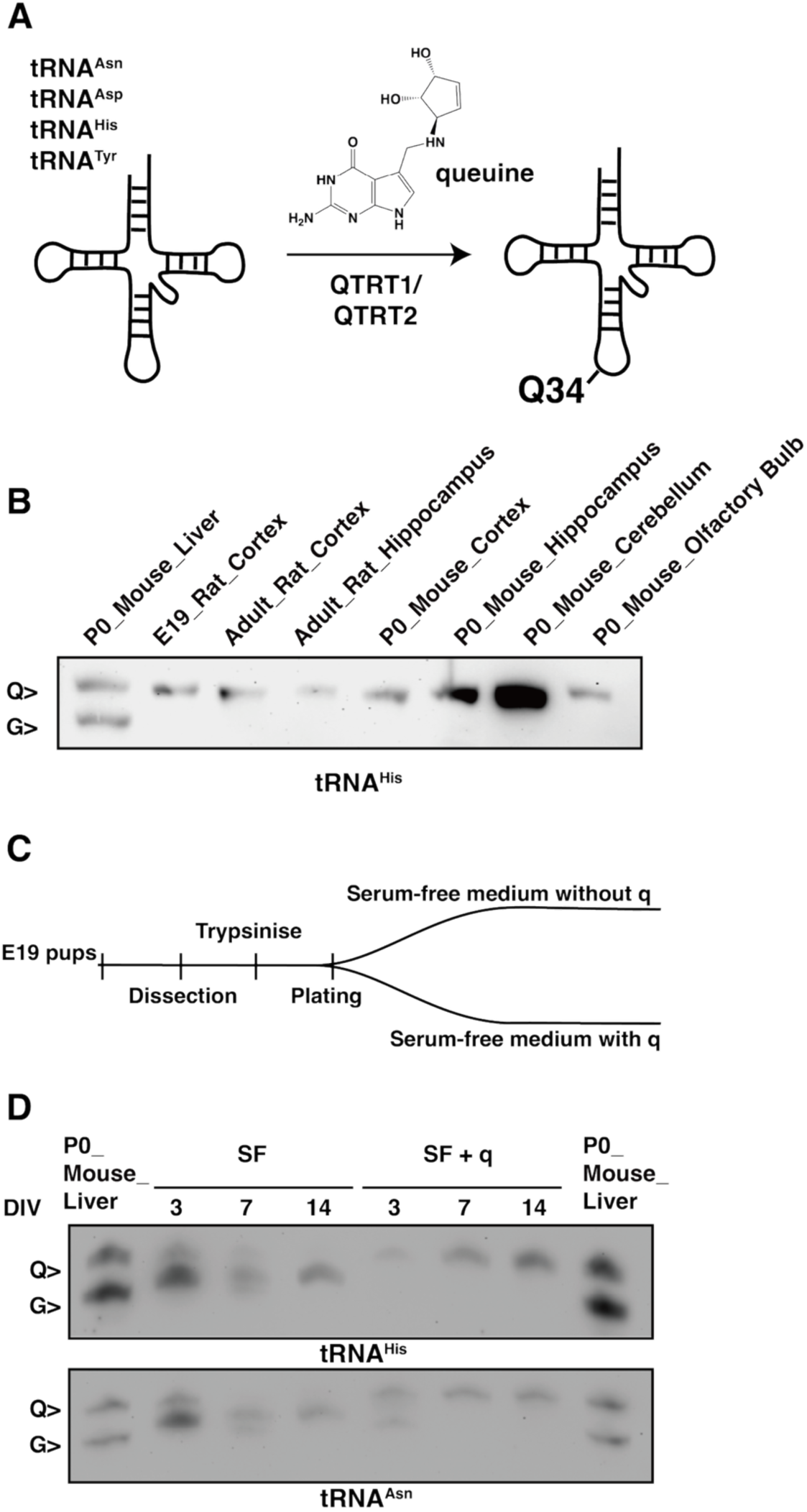
Queuosine modification dynamics in murine and primary rat culture. (A) Schematic of queuosine modification of tRNAs in eukaryotes. Eukaryotes obtain the nucleobase q from dietary sources and gut microbes. (B) Queuosine modification levels of tRNA^His^ in rat and mouse brain regions, as determined by APB gel electrophoresis followed by Northern blotting. P0 liver tissue was loaded as a marker to indicate the positions of Q- and G-tRNA. (C) Experimental setup for obtaining q-free and fully Q-modified primary rat neuronal cultures. (D) Queuosine modification levels of tRNA^His^ and tRNA^Asn^ in primary neuronal cultures over time. Analysis was performed as in (B). Data information: P0, postnatal day 0; E19, embryonic day 19; adult, mouse older than one year old; G, guanine tRNA; Q, Queuosine-modified tRNA; q, queuine; DIV, days *in vitro*; SF, serum-free.

Q modification of tRNAs shapes decoding by biasing codon-anticodon interactions, thereby fine-tuning translation to optimize both the rate and fidelity of protein synthesis (Muller *et al*, 2019). In murine and human cells, Q-tRNA favors U-ending Q codons and slows down decoding efficiency most at C-ending cognate codons (Cirzi *et al*, 2023; Tuorto *et al*, 2018; Zhao *et al*., 2023). Accordingly, transcripts with a wealth or dearth of C- versus U-ending Q codons, so-called Modification-Tunable Transcripts (MoTTs), are hypothesized to be particularly sensitive to the levels of Q modification.

Even though Q is an evolutionarily ancient tRNA modification, the absence of TGT, which results in the loss of Q modification, causes relatively mild or no phenotypes. These include defects in mitochondrial translation, glycolysis and virulence in microorganisms (Ehrenhofer-Murray, 2025). In HeLa cells, q depletion abrogates Q modification and causes increased protein aggregation, which triggers endoplasmic reticulum stress and activates the unfolded protein response (Cirzi *et al*., 2023; Tuorto *et al*., 2018; Zhao *et al*., 2023). Although, mice lacking QTRT1 are viable and fertile and have no obvious developmental defects (Cirzi *et al*., 2023), behavioral characterization revealed increased movement speed and locomotion activity as well as deficits in learning and memory, with stronger effects in females. Furthermore, Qtrt1-deficient mice showed reduced neuronal density in the dentate gyrus and CA1 regions of the hippocampus, as well as reduced length and arborization of granular pyramidal neurons. Whether this is related to a Q-mediated protein folding defect is not known. Q modification thus may be particularly important in neurons, where the precise control of protein synthesis is essential, as it supports codon-biased translational accuracy and efficiency required for neurite outgrowth, synaptic function, plasticity and longevity of the neurons (Buffington *et al*, 2014; Holt & Schuman, 2013; Pfeiffer & Huber, 2006; van Kesteren *et al*, 2006). In addition, Q modification may enhance neuronal resilience by stabilizing tRNAs under stress and supporting mitochondrial homeostasis (Wang *et al*, 2018), processes that are especially critical in neurons given their high metabolic demand and sensitivity to oxidative stress (Cobley *et al*, 2018; Harris *et al*, 2012; Lennie, 2003).

Queuine has been proposed to exert neuroprotective effects. In *in vitro* models of Parkinson’s and Alzheimer’s disease, q supplementation reduced pathological α-synuclein and tau phosphorylation, helped dopaminergic neurons survive, preserved neurite structure, and lowered TNF-α in a model for chronic Aβ, which together suggest that q may protect neurons by improving proteostasis and reducing stress-related damage (Richard *et al*, 2021). Another *in vivo* study showed that blocking the eukaryotic TGT pathway with an artificial TGT substrate could drive disease remission in a model for chronic multiple sclerosis, further linking Q modification to neuroinflammatory disease (Varghese *et al*, 2017).

Q deficiency in principle can be achieved by raising animals or culturing cells in the absence of Q and q. However, this is technically challenging. For instance, mice must be devoid of gut microbes (i.e. be axenic) and must be fed a synthetic, q-free diet, which is poorly tolerated (Tuorto *et al*., 2018). Also, cultured cells are commonly grown in serum-containing media, which contains high levels of both Q and q. Consequently, Q-free conditions require cultivation in a defined synthetic growth medium lacking Q or q. Alternatively, animals or cells mutated for QTRT1 or QTRT2 can be employed, since the absence of TGT prevents Q modification of tRNAs. However, in this case, it is possible that non-enzymatic functions of TGT could contribute to the observed phenotypes. Furthermore, the generation of TGT-mutant mice requires backcrossing, which may permit compensatory mechanisms to emerge (Rashad *et al*, 2024; Tuorto *et al*., 2018).

To overcome the limitations of genetic manipulations and directly assess the effect of Q modification on neuronal development, survival, morphology and activity, we use primary cortical neurons grown in synthetic medium lacking Q and q. Using this model system, we show that although the lack of Q modification did not affect neuronal survival and the ratio of excitatory vs. inhibitory neurons, Q modification is needed for proper neuronal arborization and density of inhibitory synapses. In the absence of Q modification, we observed enhanced inhibitory innervation of excitatory neurons which resulted in inhibition of calcium transients in cortical neurons. Altogether, our results show that tRNA Q modification directly influences neuronal structure and suggest that Q modification is critical for proper neuronal network function.

## Methods and materials

### Animals

Wistar Unilever HsdCpb:WU (Envigo) rats were obtained from the animal facility of the University Medical Center Hamburg-Eppendorf, UKE, Hamburg, Germany, from the Leibniz Institute for Neurobiology, Magdeburg, Germany and from Neuroscience Research Center / Charité CrossOver, Berlin, Germany. Protocols for preparing primary neuronal cultures were approved by the local authorities of the city-state Berlin (Landesamt für Gesundheit und Soziales) as well as the local authorities of the city-state Hamburg (Behörde für Gesundheit und Verbraucherschutz, Fachbereich Veterinärwesen), C57BL6/J mouse at postnatal 0 (P0) were used in this study. Mice were obtained from Janvier laboratories and bred at the animal facility of Humboldt-Universität zu Berlin.

### Preparation of primary cortical neuronal cultures and brain tissues

Primary rat neurons were prepared as described previously (Kapitein *et al*, 2010) with some modifications. Briefly, cortices were extracted from E18 rat embryos and treated with 0.25% trypsin for 15 min at 37 °C. Subsequently, the tissues were physically dissociated by pipetting through a 26G needle and filtered to remove large clumps. The cell number in the suspensions was counted with a Neubauer chamber, and cells were plated on poly-L-lysine-coated 18-mm glass coverslips or 10-cm petri dish at the desired density of 20,000 cells (extra low density) or 40,000 cells (low density) or 60,000 cells (high density) per 1 mL or 2.5 million cells per 10 mL. To prevent q uptake from the medium, NeuroCult™ neuronal plating medium (STEMCELL Technologies Catalog#05713), a serum-free medium, supplemented with 200 mM L-Glutamine and 2 mg/mL L-Glutamic acid was used to dissociate and plate neurons. After 1 hour, the plating medium was replaced with BrainPhys neuronal medium supplemented with SM1 (STEMCELL Modified-1) and 0.5 mM glutamine. Cells were grown at 37 °C with 5% CO_2_ and 95% humidity. Primary cell cultures were fed weekly by adding 20% of fresh medium. For all q-supplemented conditions, 40 nM of queuine (kindly provided by Hans-Dieter Gerber and Gerhard Klebe, Universität Marburg (Gerber & Klebe, 2012)) was added after replacing the plating medium with BrainPhys neuronal medium.

Mouse tissues were isolated from the sacrificed mice, washed with PBS and snap-frozen in liquid nitrogen. Subsequently, TRIzol (Invitrogen) was added to proceed with RNA isolation.

### RNA isolation

50-100 mg of tissue or 10^5^-10^7^ cells were homogenized in 1 mL of TRIzol (Invitrogen), and small RNAs were extracted according to the manufacturer’s protocol (PureLink™ miRNA Isolation Kit (Invitrogen)) with slight adjustments. The samples suspended in TRIzol were vortexed for 2 min after adding 200 μL chloroform and glass beads, and then centrifuged at 16 000 rcf, 4 °C for 15 min. After adding 215 μL ethanol to the upper phase, the samples were transferred to a spin cartridge and centrifuged at 12 000 rcf for 1 min. 700 μL of ethanol was added to the flow-through, and the samples were transferred to a new spin cartridge followed by centrifugation at 12 000 rcf for 1 min. After washing the cartridge with wash buffer, small RNAs were eluted with 30 μL of DEPC-treated water. Small RNA concentrations and purity were analyzed on the NanoDrop One. Small RNA integrity was assessed by 12% 7 M urea PAGE. Samples were stored at -20 °C.

### Acryloyl aminophenyl boronic acid (APB) gel and Northern Blot

µg small RNA of each sample was deacylated with 100 nM Tris-HCl (pH 9.0) at 37 °C for 30 min. 2X RNA loading dye (New England Biolabs) was then added to each sample, and denatured at 70 °C for 5 min. 15% 7 M urea gels with 5 mg/mL 3-(acrylamido)-phenylboronic acid (APB) were used to separate the small RNA as previously described (Igloi & Kossel, 1985). Electrophoresis was performed in 1X TBE buffer for approximately 2 h at 30 mA. The RNA was transferred onto a Biodyne B Nylon membrane (0.45 μm) in 0.5X TBE at 150 mA, 4 °C for 90 min. The membrane was then UV cross-linked and baked for 30 min at 60 °C. Pre-hybridization was performed with DIG Easy Hyb buffer (Roche) at 60 °C for at least 30 min, and the hybridization was performed at 60 °C overnight using a denatured biotinylated probe with a final concentration of 30 ng/mL in North2South™ Hybridization Buffer (Thermo Fisher Scientific). The next day, the membrane was washed 3 times at 60 °C for 20 min using the North2South™ Hybridization Stringency Wash Buffer (Thermo Fisher Scientific). tRNA was detected using the Chemiluminescent Nucleic Acid Detection Module Kit (Thermo Fisher Scientific). Briefly, the membrane was incubated with the blocking buffer, Streptavidin-HRP was added after 15 min at a dilution of 1:300 and blocked for another 15 min. The membrane was transferred to a new container and washed 4 times with wash buffer for 5 min with agitation. The membrane was placed in Substrate Equilibration Buffer for 5 min before developing the blot with Chemiluminescent Substrate Working Solution. The blot was imaged using the ChemiDoc Imaging system (Bio-Rad).

The oligonucleotide probe sequences were:

tRNA^His^: 5′-Biotin-TTG CTG CGG CCA CAA CGC AGA G-3′

tRNA^Asn^: 5′-Biotin-CCT TTC GGT TAA CAG CCG AAC GCG C -3′

tRNA^Asp^: 5′-Biotin-TCT CCC GCG TGA CAG GCG GGG ATA -3′

tRNA^Tyr^: 5′-Biotin-ACC TAA GGA TCT ACA GTC CTC CGC TC-3′

After hybridization and detection, the membrane was stripped and probed with a new probe as previously described (Cirzi & Tuorto, 2021). Briefly, the membrane was incubated in stripping buffer (50 mL formamide, 5 mL of 1 M Tris-HCl (pH 7.5), 5 g of SDS and up to 100 mL H_2_O) at 80 °C for 2 h. Then, the membrane was neutralized in 2X SSC (Saline-Sodium Citrate) for 15 min. The membrane was then ready to be probed with a new probe.

### Immunocytochemistry (ICC)

For immunocytochemistry, cells were fixed with 4% paraformaldehyde/ 4% sucrose in PBS for 10 minutes at room temperature (RT) and washed 3 times with PBS. The cells were then permeabilized (0.2%Triton X-100 in PBS) for 10 minutes, washed 3x with PBS and blocked with blocking buffer (0.1% TritonX, 10% heat-inactivated horse serum (Gibco), in PBS) for one hour at RT. Primary antibodies were added in blocking buffer overnight at 4 °C. Cells were washed again with PBS three times before secondary antibodies were added in blocking buffer for 1 hour at RT. Finally, the coverslips were washed three times with PBS and then mounted in Mowiol on microscope slides before imaging. Detailed information about the antibodies used in this study can be found in Supplemental table 1.

### Fixed cell imaging: spinning disc confocal microscopy

Spinning-disc confocal microscopy was performed on a pco.edge 4.2 bi sCMOS camera (Excelitas PCO GmbH) with a Nikon Eclipse Ti-E controlled by VisiView software. Corresponding filters were used for acquisition. Multichannel z-stacks were taken sequentially. Excitation lasers used were 405 nm, 488 nm, 561 nm and 640 nm. Fixed imaging was performed using 20X objective (Nikon, Plan Fluor 20X/0.50), 40X objective (Nikon, Plan Fluor 40X/1.30 oil), 60X oil objective (Nikon, ApoTIRF 60X/1.49) and 100X oil objective (Nikon, ApoTIRF 100×/1.49 oil). The final pixel size was 65 nm. For analysis of cell populations and cell proportion experiments, tile scans of a maximum 2 × 2 mm areas were performed to acquire large areas. Tile scan images were stitched in FIJI.

### Calcium imaging

For calcium imaging experiments, primary rat cortical neurons were infected with a pAAV-syn-ruby-P2A-GCaMP6s virus (final concentration: 3.76E+13vg/mL) at day *in vitro* (DIV) 9 and imaged at DIV 14. The samples were incubated in a stage top incubator (Okolab) at 37 °C, 5% CO_2_ and 90% humidity atmosphere. Neurons were imaged in artificial cerebrospinal fluid (aCSF) (125 mM NaCl, 25 mm NaHCO₃, 10 mm Glucose, 2.5 mM KCl, 1.25 mM NaH₂PO₄, 2.0 mM CaCl₂) without magnesium. Images were acquired at 10 frames per second for 2 min.

### Data analysis and statistics

The experimental groups were blinded prior to data analysis. All analysis was done on raw images. All analyses involving intensity measurements were performed using the same analysis parameters for both groups.

### Determination of cell apoptosis

Neurons (10,000-20,000 cells/mL) were fixed at DIV 3, 4, 6, 8, 10, 12 with 4% PFA/4% sucrose in PBS. Cleaved-caspase-3 antibody was used to determine the cells undergoing apoptosis, and MAP2 antibody was used to label neurons. DAPI staining of DNA was used to visualize the nuclei. After stitching of tile scan images, caspase-3 and MAP2 positive cells were quantified by first applying a threshold and then counted using AnalyzeParticle in Fiji (Schindelin *et al*, 2012). The number of caspase-3 positive cells was divided by the number of MAP2 positive cells to calculate the percentage of cells undergoing apoptosis. Individual density of MAP2 and caspase-3 are calculated by dividing the absolute number of MAP2 positive cells and caspase-3 positive cells by the imaged area.

### Analysis of inhibitory neuron density

DIV14 neurons (20,000-30,000 cells/mL) were fixed at RT with 4% PFA/4% sucrose in PBS for 10 min. The MAP2 and GAD1 antibodies were used to visualize somato-dendritic morphology and GABAergic neurons. Cell bodies were selected with circular regions of interest (ROIs) in the MAP2 channel, and the mean intensity was measured in both channels. A threshold was determined to select neurons positive for GAD1, and the number of positive neurons was divided by the total number of MAP2-positive neurons. For GAD1, somata were additionally detected in the GAD1 channel due to often lower MAP2 presence in the soma. Individual density of MAP2 and GAD1 was calculated by dividing the absolute number of MAP2 positive cells and GAD1 positive cells by the imaged area.

### Analysis of dendritic morphology

Dissociated neurons were fixed at DIV 3, 7, 14 (40,000 cells/mL) with 4% PFA/ 4% sucrose in PBS. Antibodies against MAP2 were used to visualize the cell body and dendrites. DAPI staining was additionally used to visualize the nucleus. The analysis of dendritic arborization was performed using the Fiji plugin Neuroanatomy. The paths were determined by manually clicking the starting point and the ending point of a dendrite. After the whole cell was skeletonized, the number of branches as well as the length of branches were measured. For Sholl analysis, the concentric circles were set with a radius step of μm, and the number of intersections was automatically counted according to the skeletonized paths.

### Analysis of synaptic density

Low-density DIV14 neurons (40,000 cells/mL) were fixed for 10 min at room temperature. For inhibitory and excitatory synapse stainings, fixed neurons were stained either with antibodies against VGAT and gephyrin for pre- and postsynaptic sites of inhibitory synapses, or with antibodies against Shank3 and bassoon to visualize the pre- and postsynaptic compartments of excitatory synapses. MAP2 antibody was used to visualize cell bodies and dendrites. All samples were imaged using a spinning disk confocal microscope (Visitron Systems) and processed in Fiji.

To quantify excitatory synapses, secondary dendrites or distal dendrites were imaged for MAP2, Shank3 and bassoon. After creating composite images, colocalizing signals were counted with multipoint tool and normalized to the length of measured dendrite.

To quantify inhibitory synapses, images of MAP2, VGAT and gephyrin were merged to create composite images. The analysis of inhibitory synapses was performed using the Fiji plugin Comdet. ROIs were defined using the polygon selections tool to outline the dendritic branches. Next, we restricted the maximum distance between the center of co-localized gephyrin and VGAT particle to 5 × 5 pixels (x and y). To calculate density, the number of co-localized gephyrin-VGAT particles was divided by the length of the measured dendrite.

To calculate the excitatory-to-inhibitory (E/I) synaptic ratio, the mean numbers of excitatory and inhibitory synapses per 10 µm were first summed to obtain the total synapse density per 10 µm. The mean number of excitatory or inhibitory synapses was then divided by this total to determine the relative proportion of each synapse type, expressed as a percentage.

### Statistics and image presentation

Statistical analysis was performed in GraphPad Prism (version 9.1), with detailed specifications of tests, significance levels, n numbers, and biological replicates as included in the figure legends. Data are represented as mean ± SEM unless otherwise stated. Before parametric statistical analysis, we used the one-sample Kolmogorov-Smirnov and D’Agostino & Pearson test to assess the normal distribution of our datasets. For comparisons between two groups, normally distributed data were analyzed using paired or unpaired Student’s t-tests, as appropriate, whereas non-normally distributed unpaired data were analyzed using the Mann-Whitney test.

Individual channels in multicolor images are contrasted for better representation. No other modifications were done unless otherwise stated.

## Results

### Q modification of tRNAs is gradually lost in primary cortical neurons cultured in q-free medium

The Q modification level of tRNAs is highly dynamic during fly development (Zaborske *et al*, 2014). In mice, Q modification levels vary across tissues, with particularly high levels reported in the brain of 3-month-old animals compared to other tissues (Cirzi *et al*., 2023). We therefore first investigated Q modification levels in different murine brain regions, as well as in brain tissue from embryonic and newborn rat and mice where the only source of Q is maternal. Q modification levels can be measured by polyacrylamide gel electrophoresis containing 3-(acrylamido)-phenylboronic acid (APB), which causes slower migration of Q- compared to the non-modified tRNAs (Igloi & Kossel, 1985). Interestingly, Northern blot analysis of APB gels for tRNA^His^ and tRNA^Asn^ showed that these tRNAs were fully Q-modified in all examined mouse and rat brain tissues, which may be the consequence of the low neuronal cell turnover rate (Figure 1B, Suppl. Fig. 1A).

Next, to establish primary rat cortical neuron cultures, we used E19 rat cortex as starting material and developed a serum-free (SF) culture system to restrict serum-derived sources of Q and q (Figure 1C). This system enables controlled q supplementation, thus allowing the comparison of neuronal morphology and activity under Q-depleted (SF) and Q-modified (SF + q) conditions. Indeed, after 3 days of *in vitro* in the absence of q, only a minor proportion of tRNA^His^ and tRNA^Asn^ in neurons retained Q modification, and at the later timepoints DIV7 and DIV14, they were mostly unmodified (Figure 1D). Conversely, supplementation of q in the growth medium supported full Q modification in the neuronal cells as early as DIV3 (Figure 1D). Q modification levels of tRNA^Asp^ and tRNA^Tyr^ could not be assessed, as their subsequent conversion to manQ and galQ, respectively, prevents identification of the corresponding Q-modified forms by APB-gel Northern blotting (Suppl. Figure 1B).

### Alteration of Q modification levels does not impact neuronal survival and subtype composition

The absence of Q modification has previously been reported to lead to a decrease of neuronal density in the dentate gyrus of the mouse brain (Cirzi *et al*., 2023). Whether the absence of Q modification directly induces programmed cell death in neuronal cells or plays a role in neuronal development was not determined. To investigate the relationship between Q modification and cell death, we quantified apoptosis rates in neurons with and without q supplementation. During apoptosis, procaspase-3 is cleaved by upstream caspases like caspase-8 or caspase-9, and the cleaved caspase-3 will subsequently execute apoptosis in the cell (Porter & Janicke, 1999). We therefore quantified cleaved-caspase-3-positive neurons from DIV3 to DIV12 to calculate the rate of neuronal cell death (Figure 2A, 2B). Although neurons cultured under SF conditions showed higher apoptosis at earlier DIVs, whereas SF + q cells showed higher values at later DIVs, no significant difference in the overall apoptosis time course was observed between the two groups. Additionally, we quantified MAP2-positive neurons, which confirmed that both groups of neurons exhibited similar cell density (Figure 2D). To gain further information of neuronal viability, we also plotted the total area under the curve of the cell death curve and cell density (Figure 2C & 2E). This showed that primary neurons with and without q supplementation exhibited similar survival rates.

**Figure 2.**
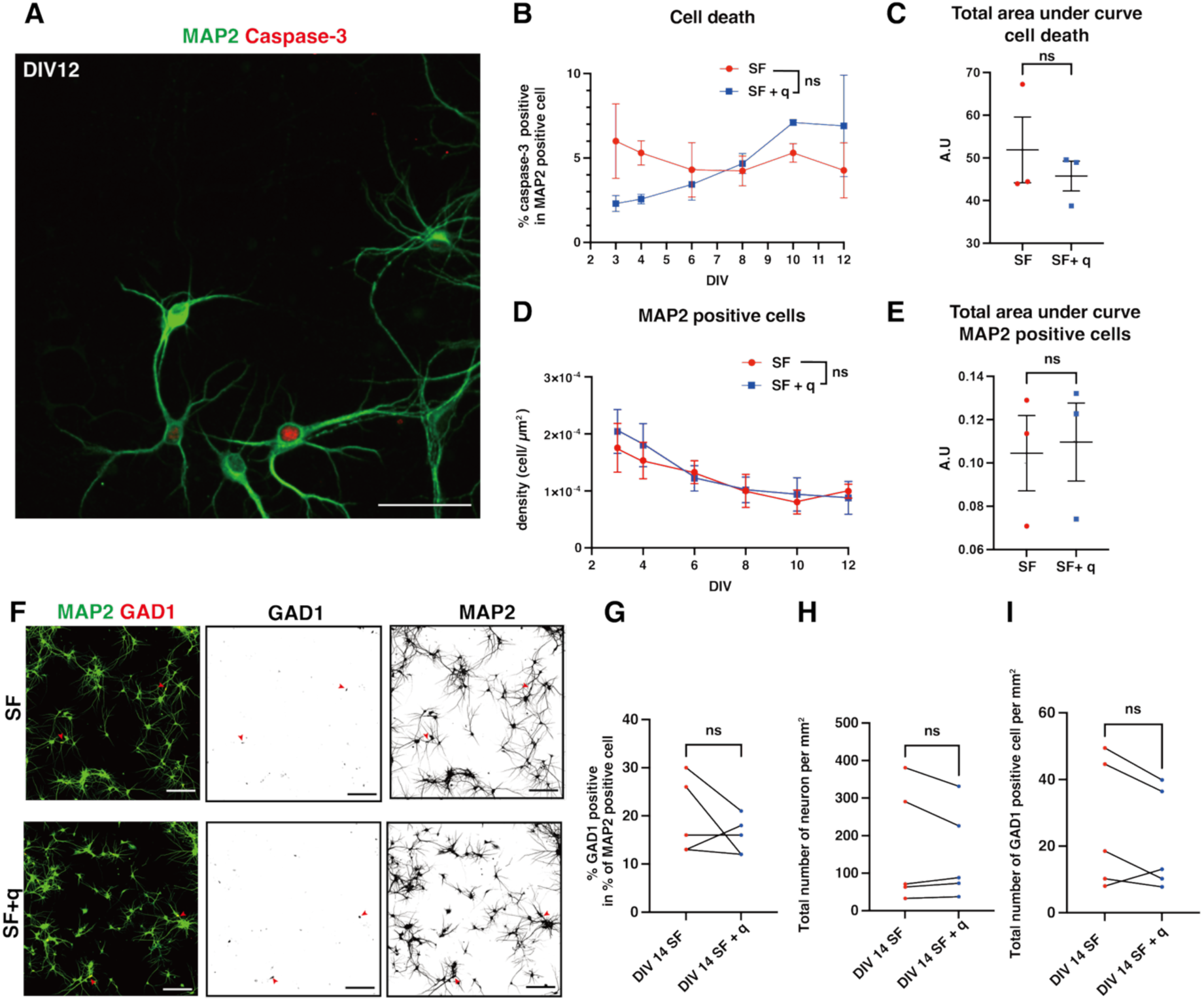
Queuosine modification did not affect apoptosis and the proportion of cell types in primary cortical neurons. (A) Representative image of DIV12 primary cortical neurons stained with the apoptosis marker Caspase-3 and the neuronal marker MAP2. Scale bar is 50 µm. (B) Quantification of cleaved-caspase-3-positive neurons in primary cortical cultures with or without q supplementation at DIV3, 4, 6, 8, 10, and 12. n= 3 coverslips from 3 independent replicates for each time point. Two-way ANOVA showed no significant difference between the two groups. ns, not significant. (C) Area under curve analysis of the apoptosis time course shown in (B). Welch’s t test (two tailed), ns for not significant. (D) Quantification of the density of live neurons. n= 3 coverslips from 3 independent replicates for each time points. Two-way ANOVA showed no significant difference between the two groups. ns, not significant. (E) Area under curve analysis of the cell density curve shown in (D). Welch’s t test (two tailed), ns for not significant. (F) Representative images of excitatory and inhibitory neurons at DIV14. Scale bar is 150 µm. (G) Relative amount of inhibitory neuron at DIV14 under SF and SF + q conditions. n= 5 coverslips from 3 individual experiments. (H) Absolute density of all neurons quantified in (G). Welch’s t test (two tailed), ns for not significant. (I) Absolute density of inhibitory neurons quantified in (G). Welch’s t test (two tailed), ns for not significant. All data are presented as mean ± SEM.

Excitatory and inhibitory neurons derive from different precursors and follow different developmental timelines (Molyneaux *et al*, 2007; Wonders & Anderson, 2006). It thus was conceivable that some cell types would be more sensitive to the presence or absence of Q than others. We next examined the proportion of different neuronal subtypes in the culture. Alterations during early neuronal development could selectively disrupt the inhibitory GABAergic system, thereby alter the excitatory-inhibitory balance and promote pathological neuronal hyperexcitability (Nelson & Valakh, 2015). We therefore asked whether q supplementation during *in vitro* culturing alters the number of inhibitory neurons.

In primary neuronal culture, glutamatergic and GABAergic neurons are likely to mature around DIV14 (Bjorklund *et al*, 2010). We quantified the relative number of inhibitory neurons at DIV14 using GAD1 as a marker for inhibitory neurons and MAP2 as a general neuronal marker (Figure 2F). Both SF and SF + q neurons exhibited an inhibitory neuron proportion of approximately 10-30% that did not differ between the two experimental conditions (Figure 2G). Furthermore, q supplementation did not alter the absolute density of inhibitory neurons or of the overall neuronal population (Figure 2H & 2I).

### Q modification promotes dendritic arborization in primary cortical cultures

Neurons are highly polarized cells, and their growth and differentiation strongly depend on protein synthesis. Since Q affects the translational speed, we next asked if q supplementation influences dendritic branching and growth. We thus measured neuronal arborization by performing Sholl analysis. At DIV3, there were no differences between SF and SF + q conditions (Figure 3A). Of note, at this early stage, the maternally supplied Q modification is still present (Figure 1D).

**Figure 3.**
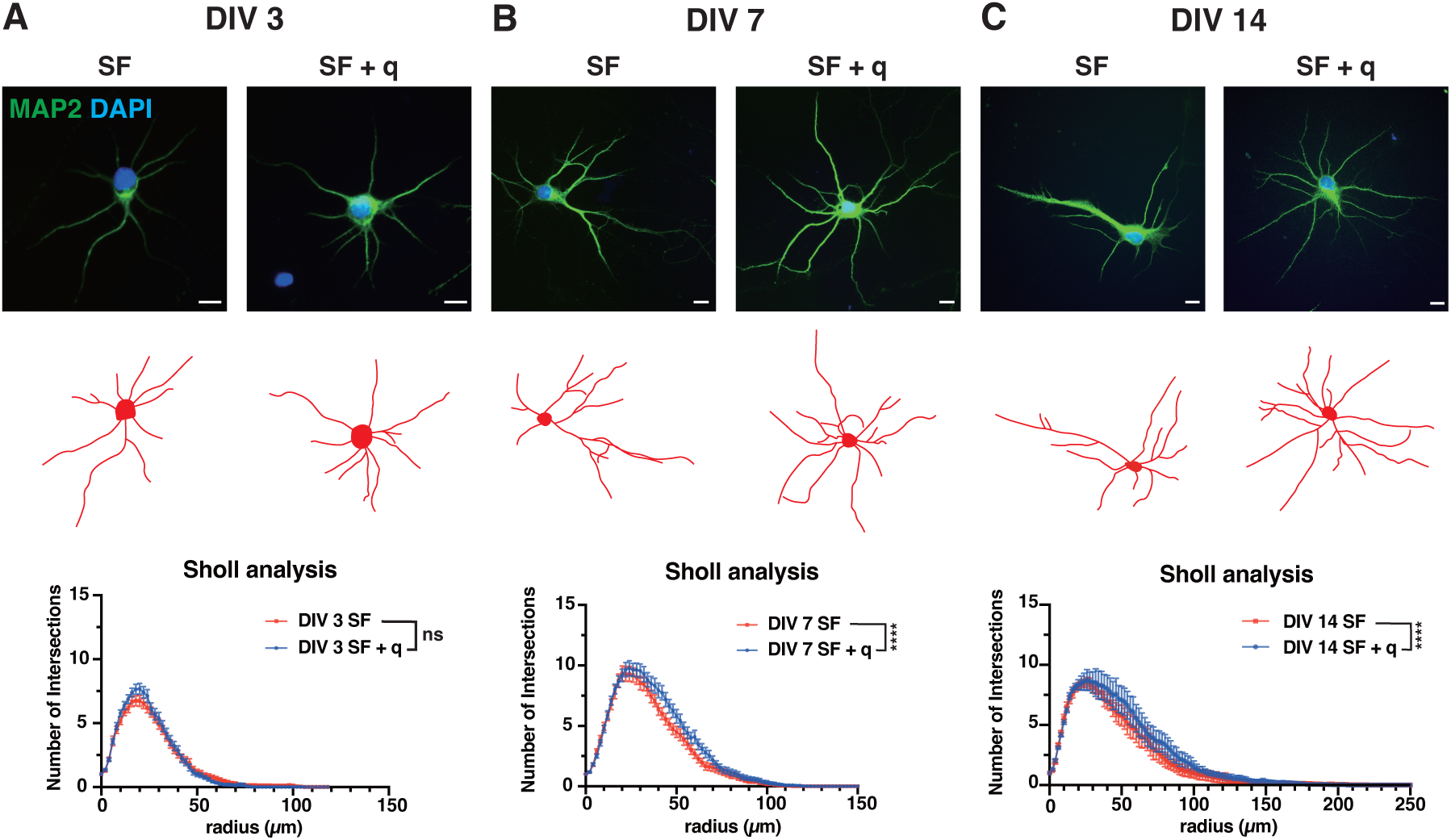
Q modification causes increased neuronal arborization. (A) Sholl analysis of Q-modified (SF + q) and non-Q-modified (SF) primary neurons at DIV3. n(SF)= 51 cells; n(SF + q) = 56 cells, from 3 independent biological replicates. Two-way ANOVA showed no significant difference between the two groups. ns, not significant. (B) Sholl analysis of primary neurons cultured under SF + q and SF conditions at DIV7. n(SF) = 49 cells; n(SF + q) = 43 cells, from 3 independent biological replicates. Two-way ANOVA revealed a significant difference between the two groups. **** for p < 0.0001. (C) Sholl analysis of primary neurons cultured under SF + q and SF conditions at DIV14. n(SF) = 48 cells; n(SF + q) = 57 cells, from 3 independent replicates. Two-way ANOVA revealed significant differences between the two groups. **** for p < 0.0001. Scale bar is 10 µm. Values are presented as mean ± SEM.

Next, we performed investigation of neuronal arborization at later stages. At DIV7 and DIV14, when neurite outgrowth is more advanced and Q modification in neurons cultured under SF conditions is fully depleted. This allows comparison between Q-modified versus unmodified conditions (Figure 3B & 3C). Sholl analysis showed a rightward shift of the SF + q curve both at DIV7 and DIV14, indicating greater arbor complexity under Q-modified conditions. The similar pattern observed at both time points suggests that the loss of Q modification does not progressively worsen the dendritic simplification, but may instead cause a sustained delay in neurite outgrowth.

### Q modification controls inhibitory but not excitatory synapse density

We next aimed to investigate the impact of Q modification on synapse density in mature cortical neurons (DIV14). For this, Shank3, a postsynaptic excitatory marker (Wan *et al*, 2021), and bassoon, a presynaptic marker, were used to assess excitatory synapses (Figure 4A). We counted excitatory synapse density on the secondary dendrites and found no significant difference between conditions with or without Q modification (Figure 4B).

**Figure 4.**
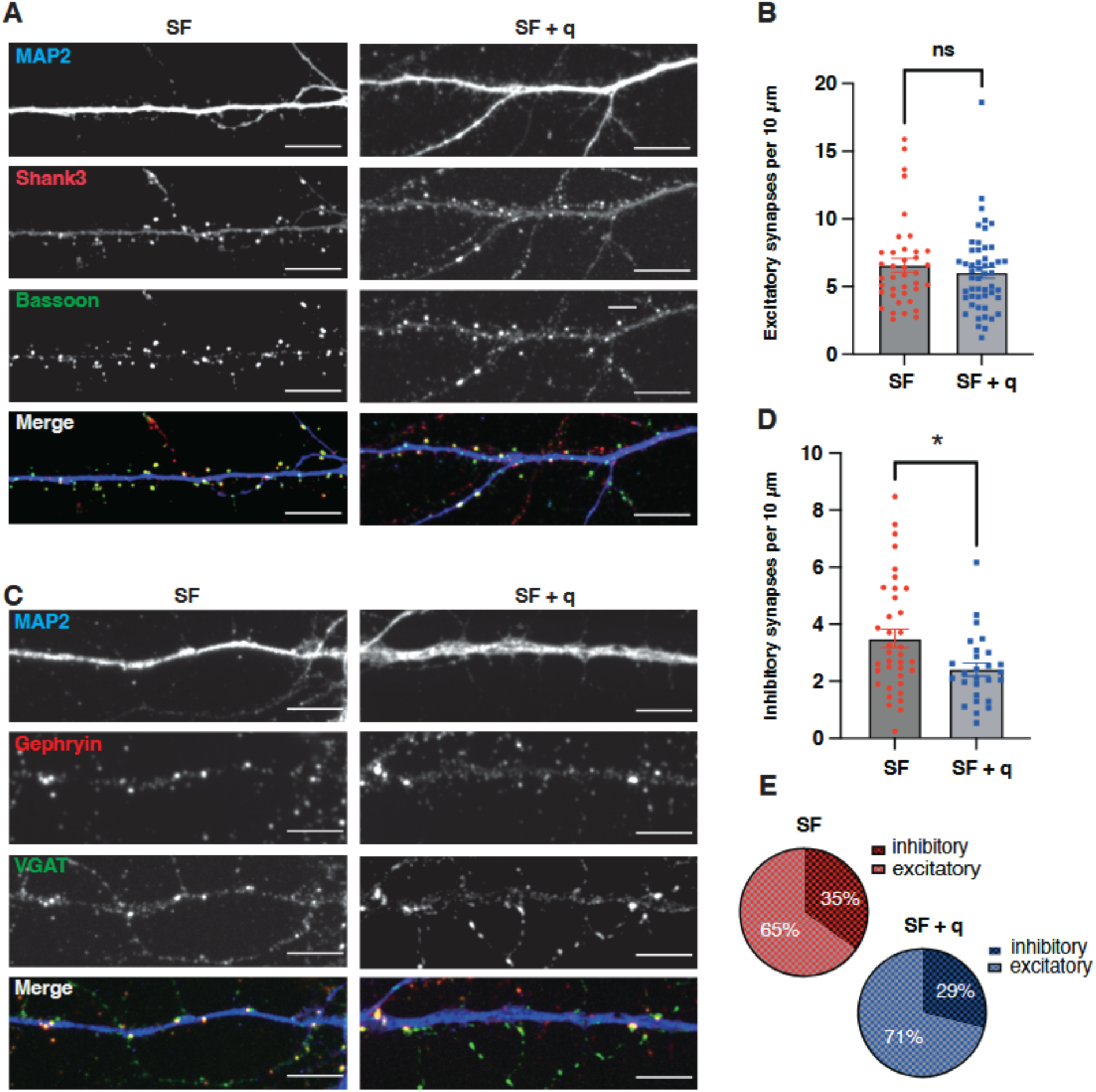
Queuosine modification reduces density of inhibitory synapses in primary cortical neurons. (A) Representative images of neurons stained with markers for excitatory synapses. Scale bar is 5 µm. (B) Quantification of excitatory synaptic density in neurons under SF + q and SF conditions. Mann-Whitney test (two-tailed), ns, not significant. Values are presented as mean ± SEM, n(SF) = 39 cells; n(SF + q) = 50 cells, from 3 independent cultures. (C) Representative images of neurons stained with markers for inhibitory synapses. Scale bar is 5 µm. (D) Quantification of the density of inhibitory synapses in Q-modified and non-Q-modified neurons. Mann-Whitney test (two-tailed), * for p < 0.05. Values are presented as mean ± SEM, n(SF) = 37 cells; n(SF+ q) = 27 cells, from 3 independent cultures. (E) Relative proportion of excitatory and inhibitory synapses in neurons cultured under SF + q and SF conditions.

We next explored the impact of Q modification on inhibitory synapses density. To this end, the density of the inhibitory pre- and postsynaptic markers, vesicular GABA transporter (VGAT) and gephyrin, respectively, was determined (Figure 4C). Importantly, in the presence of Q modification, the density of inhibitory synapses was significantly decreased (Figure 4D). In SF cultures, neurons carried approx. 3.5 inhibitory synapses per 10 µm dendrite, whereas the density was decreased to 2.4 inhibitory synapses per 10 µm in SF + q cultures, corresponding to a 31.4% decrease. This indicates that Q modification does not influence excitatory connectivity but limits the inhibitory innervation. These morphological features suggest that, eliminating Q modification can shift the excitatory/ inhibitory synaptic ratio towards a stronger inhibition (Figure 4E).

### Q modification affects the activity of neuronal cultures

Given that the above experiments revealed an effect of Q tRNA modification on the density of inhibitory synapses, we next asked whether Q modification had an impact on spontaneous neuronal activity of cultured primary cortical neurons. Here, we employed calcium imaging to study neuronal activity in neurons with or without Q modification. Calcium imaging relies on the tight coupling between electrical activity and intracellular Ca²⁺ dynamics. We recorded calcium transients in SF and SF + q neurons by expressing GCaMP6s in neurons, and the frequency and amplitude of calcium signals were analyzed (Figure 5A).

**Figure 5.**
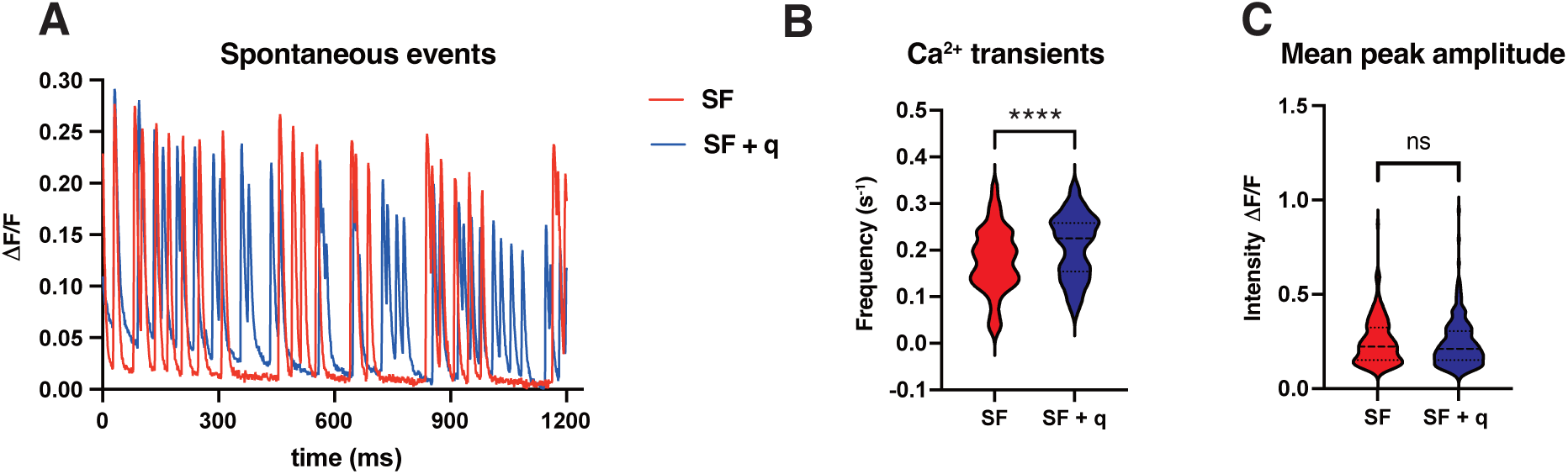
Q modification of tRNAs causes increased spontaneous calcium transients in primary cortical neurons. (A) Representative graph of calcium transient events in Q-modified (SF + q) and non-Q-modified (SF) neurons. (B) Peak frequency of calcium transients in neuron. Mann Whitney test (two tailed), **** for p <0.0001. (C) Mean peak amplitude of calcium transients in neurons. Welch’s t test (two tailed), ns for not significant. Values are represented as density estimates and summary statistics, n(SF) = 242 cells; n(SF + q) = 241 cells, from 3 independent biological replicates.

Importantly, we found that SF + q neurons fired 17% more frequently compared to SF neuron, whereas the amplitude was unaffected (Supplementary Video 1 & 2, Figure 5B & 5C). This is in good agreement with the reduced density of inhibitory synapses in SF + q neurons. The calcium transients do not scale linearly with the action potential, but amplitude and frequency may be associated distinct aspects of neuronal morphology, such as synaptic organization and dendritic computation. In summary, these results showed that Q modification enhances neuronal arborization, modulates inhibitory synapses, and is associated with spontaneous firing in primary cortical neurons.

## Discussion

tRNA modifications have emerged as important contributors to a wide range of neurological disorders (Guo *et al*, 2024). While previous studies have primarily focused on cognitive dysfunction and systemic phenotypes caused by depletion of Q modification (Cirzi *et al*., 2023; Del-Pozo-Rodriguez *et al*, 2025; Gonskikh *et al*, 2025), their impact on neuronal morphology and neuronal activity remains poorly understood. Here, in a reductionist approach using primary rat cortical neurons as a model system, we explored the impact of Q modification on these processes and its consequences for neuronal development and synaptic organization.

Queuine has been termed a longevity vitamin (Ames, 2018). The broad evolutionary conservation of Q-tRNA modification implies an important biological function that may be especially relevant for long-lived cells with intensive metabolism and protein turnover. We found that the morphology of primary cultured cortical neurons was less complex when cells were depleted for Q modification. The differences in neuronal arborization were observed at DIV7 and DIV14, suggesting that depleting Q modification constantly slows down the branching process, which depends on protein synthesis. Abnormalities in dendritic branching are tightly linked to pathological disorders such as Autism spectrum disorder and Fragile X syndrome (Kulkarni & Firestein, 2012). QTRT1 mutations have been reported to cause mild neurodevelopmental defects in mice (Cirzi *et al*., 2023). Moreover, glycosylated Q modification has been shown to be vital for post-embryonic growth in zebrafish (Zhao *et al*., 2023), suggesting the developmental role of Q modification. Yet, direct evidence that Q modification affects neurodevelopmental disorders is still rather limited. It would therefore be of interest to investigate the Q modification levels in patients with neurodevelopmental disorders and to examine whether the administration of q has beneficial effects. As gut-derived Q and q can both reach the central nervous system, they may hold therapeutic potential to help maintain neuronal health.

Another aspect of Q modification importance has been shown in case of autoimmune diseases and aging. Thus, remission of disease symptoms in the mouse model of multiple sclerosis when supplemented q implies a role for q in neuroinflammation (Varghese *et al*., 2017). A recent study showed that increase activation of the mTOR pathway in aging mice can be reversed by q (Gong *et al*, 2026). Mechanistically, balancing the Q modification levels in old mice suppresses p16/p21-driven senescence programs, re-establishes proteostasis and delay a cellular and organismal aging across different species. Together, these studies suggested therapeutic potential for q.

We furthermore demonstrated that neurons maintain the density of excitatory synapses but show a significantly lower density of inhibitory synapses in the presence of Q modification. Synapse density and distribution is mainly regulated via synaptogenesis and synapse pruning, where the former promotes synaptic connection and the latter eliminates the unused synapses (Paolicelli *et al*, 2011; Prange *et al*, 2004; Stevens *et al*, 2007). In our study, the mechanism involved in our findings is unconfirmed, but is consistent with an earlier study reporting the dysregulation of synaptogenesis in QTRT1-deficient mice (Cirzi *et al*., 2023). Q-tRNA modification in neurons might regulate neuronal circuits by controlling the number of inhibitory synapses. The excitatory/ inhibitory synapse ratio in primary cortical culture is well documented, and increased excitation/inhibition of neuronal networks result in several pathological conditions such as epilepsy and seizure susceptibility (Fattorini *et al*, 2015; le Feber *et al*, 2018; Sarnat & Flores-Sarnat, 2021). The different excitatory/ inhibitory synapse ratios observed here provide an unanticipated aspect of the regulatory effect of Q modification on synapse formation and/or synapse pruning, a finding that requires further clarification in future studies.

Interestingly, our results revealed a pronounced decrease in neuronal firing upon depletion of Q-tRNA modification, likely resulting from the observed dendritic simplification and the higher number of inhibitory synapses innervation. A more complex dendritic arbor generally implies increased connectivity, as dendrites are the primary structures responsible for receiving and integrating synaptic input (Gao, 2007; Lanoue & Cooper, 2019; Ma *et al*, 2008). In addition, the increased density of inhibitory synapses, which exert strong inhibitory control, may further contribute to neuronal dysfunction upon loss of Q modification.

In contrast to an earlier study that reported lower neuronal density in mice carrying a mutation in QTRT1(Cirzi *et al*., 2023), we observed no significant effect of Q modification on cell death in primary cortical cultures. One possibility is that QTRT1 has additional cellular functions. Moreover, that study reported decrease neuronal density in 8-month-old mice, suggesting that neuronal cell death develops occurs over an extended period or at later stages. Further, regional specificity of the impact of Q-tRNAs in different brain regions may explain the difference to our study, since QTRT1 mutation specifically reduced neuronal density in different subregions of the hippocampus in adult mice, while we employed primary cortical cultures. We further excluded potential effects of Q modification on other specific neuronal subtypes such as interneurons (Figure 2G), which have a distinct morphology and function (Laturnus *et al*, 2020).

Neurons are cells exhibiting a high energy demand, and synaptic activity for neuronal function requires substantial supplies of ATP (Faria-Pereira & Morais, 2022; Harris *et al*., 2012). Recent studies have proposed that silencing of neuronal activity is a mechanism to overcome energy shortage in neurons (Ching *et al*, 2012; Fiskum *et al*, 2021). It therefore would be interesting to investigate whether eliminating Q modification causes changes in neuronal metabolism that lead to functional defects. In this context, investigating ATP production, mitochondrial dynamics, and synaptic transmission in Q-deficient neurons may provide mechanistic insight into how tRNA modifications contribute to neuronal adaptation to energetic constraints.

In conclusion, we found that the loss of Q modification does not alter neuronal survival and subtype composition, but that it reduces dendritic arborization. In the absence of Q, more neurons exhibit an increase in inhibitory synapses, which at the functional level led to a reduced frequency of neuronal firing. This indicates that Q modification is essential for keeping the excitation/ inhibition balance in mature neurons, and it acts mostly at the level of inhibitory synapses without impacting the number of inhibitory neurons. In a clinical context, this indicates that individuals suffering from neurodegenerative disorders as well as the elderly might benefit from dietary supplementation with q.

## Authors’ contributions

N. Yang: Conceptualization, Investigation, Data curation, Formal analysis, Writing - original draft, Writing - review & editing

Y. Sun: Supervision, Project administration, Writing - review & editing

L. Mallis: Methodology, Data curation

M. Boutonnet: Methodology, Writing - review & editing

J. Bär: Supervision, Writing - review & editing

E. Ehrenhofer-Murray: Conceptualization, Funding acquisition, Project administration, Resources, Supervision, Writing - original draft, Writing - review & editing

M. Mikhaylova: Conceptualization, Funding acquisition, Project administration, Resources, Supervision, Writing - original draft, Writing - review & editing

## Acknowledgements

This work was funded by Deutsche Forschungsgemeinschaft (DFG, German Research Foundation), the Excellence Strategy –EXC-2049–390688087, DFG FOR 5705 RP4 and the DFG FOR5228 RP4 to M.M, grants EH237/19-1 and EH237/21-1 to A.E.E.-M., and funds from IZ-LIST of Humboldt-Universität zu Berlin.

